# A theory of cerebellar learning as a spike-based reinforcement learning in continuous time and space

**DOI:** 10.1101/2024.06.23.600300

**Authors:** Rin Kuriyama, Hideyuki Yoshimura, Tadashi Yamazaki

## Abstract

The cerebellum has been considered to perform error-based supervised learning via long-term depression (LTD) at synapses between parallel fibers and Purkinje cells (PCs). Since the discovery of multiple synaptic plasticity other than LTD, recent studies have suggested that synergistic plasticity mechanisms could enhance the learning capability of the cerebellum. Indeed, we have proposed a concept of cerebellar learning as a reinforcement learning (RL) machine. However, there is still a gap between the conceptual algorithm and its detailed implementation. To close this gap, in this research, we implemented a cerebellar spiking network as an RL model in continuous time and space, based on known anatomical properties of the cerebellum. We confirmed that our model successfully learned a state value and solved the mountain car task, a simple RL benchmark. Furthermore, our model demonstrated the ability to solve the delay eyeblink conditioning task using biologically plausible internal dynamics. Our research provides a solid foundation for cerebellar RL theory that challenges the classical view of the cerebellum as primarily a supervised learning machine.

---

Theories of learning in the cerebellum have historically attracted considerable interest. One of the most widely known theories, known as the Marr-Albus-Ito model (1–3), proposes that the cerebellum acts as an error-based supervised learning (SL) machine (4) to modify output and reduce the discrepancy between actual and desired outputs. In this theory, climbing fibers (CFs) deliver error or teacher signals that can trigger long-term depression (LTD) at parallel fiber (PF) and Purkinje cell (PC) synapses (5). Since the discovery of LTD in the early 1980s (6), biological research has found many other forms of synaptic plasticity in the cerebellum (7). These findings have been extending the Marr-Albus-Ito theory gradually (8–10). Furthermore, researchers have examined the learning theory using realistic spiking network models of the cerebellum (11–17). Especially, Hausknecht et al. (13) examined the learning capability of the cerebellum through a cerebellar spiking network modeled in an SL context. The network successfully performed SL and control tasks but failed to solve RL and complex recognition tasks. Geminiani et al. (17) implemented a spiking network-based cerebellar model featuring bidirectional plasticity at PF-PC and PF-molecular layer interneurons (MLIs) synapses across multiple microzones, with several microzones more likely depression and others more likely potentiation during motor control. Their results highlight the cerebellum’s capacity for error correction and adaptive learning, particularly within the context of associative learning and motor control. Both computational studies modeled the cerebellum as a conventional associative learning machine and highlighted the cerebellar learning capability, especially in the SL context.

One of the other research directions is to adapt reinforcement learning (RL) theory (18) as an alternative to conventional SL. In an RL context, there is an “agent” and an “environment”. The agent takes an action affecting the state of the environment. In response to the action, the environment sends a “reward” to the agent which represents how good/bad the action was. The agent learns appropriate actions that maximize an expected future reward. One of the most notable differences between SL and RL is feedback information from the environment. An SL machine receives errors or teacher signals representing optimal actions, while an RL machine receives only reward signals. Swain et al. (19) reported that the cerebellum fulfills three key requirements to establish reinforcement learning: it receives sensory information about the external environment and the internal environment, selects a behavior to execute, and processes evaluative feedback on the success of that behavior. Building on this idea, Yamazaki and Lennon (20) proposed the concept that the cerebellum might perform RL. Specifically, CFs deliver reward information, PCs select actions, and MLIs evaluate the current state. Synaptic weights at PF–MLI as well as PF–PC synapses are modulated by the CF signals. The researchers interpreted this structure as an RL machine, specifically as an actor-critic model (18). However, there has been no detailed spike-based implementation for the cerebellar RL concept.

In this research, based on a continuous time actor-critic framework with spiking neurons (21), we implemented a cerebellar spiking network as an actor-critic model based on the known anatomical properties of the cerebellum. In particular, we modified a weight-updating rule of the framework for the cerebellum. We evaluated the model using a simple machine learning benchmark task, known as the mountain car task (22). We also examined the behavior of the model using delay eyeblink conditioning (23), which is a standard cerebellum-dependent motor learning task. To our knowledge, the model presented here is the first cerebellar spiking network model that can act as an RL machine.

## Results

### Spike-based implementation of cerebellum-style RL

In this research, we implemented a cerebellar spiking network model as an RL machine (Figure 1) while referring to the cerebellar RL concept (20) and the spiking actor-critic framework (21). Parallel fibers (PFs) excite stellate cells (SCs), basket cells (BCs), and Purkinje cells (PCs). SCs and BCs inhibit PCs. BCs and PCs inhibit each other, and also both have a self-inhibitory connection. More specifically, BCs and PCs are divided into multiple groups, which correspond to action choices as described in the section titled ‘Actor and action selection’. BCs inhibit PCs in the same group and BCs in other groups, whereas PCs inhibit BCs in the same group and PCs in other groups. PCs also inhibit neurons in the deep cerebellar nuclei (DCN), which are also divided into multiple groups. CFs provide excitatory inputs to SCs and PCs. We used CF signals to trigger synaptic plasticity. Therefore, we did not implement actual CF–SC and CF–PC connections.

**Figure 1.**
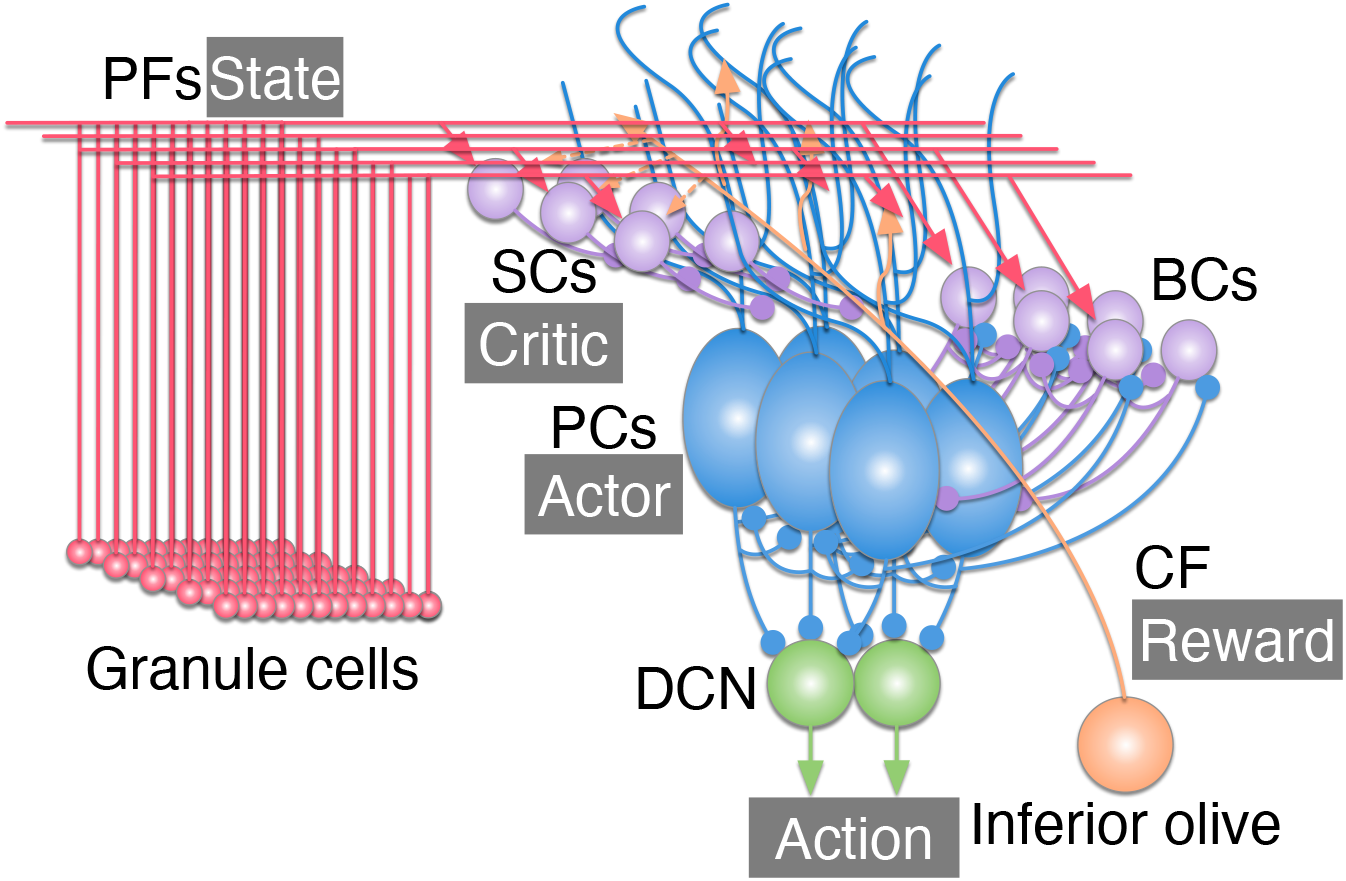
Structure of the present cerebellar network implemented as an actor-critic model. Red, purple, blue, green, and orange cells represent granule cells, molecular layer interneurons (SCs and BCs), PCs, DCN, and inferior olive, respectively. PFs deliver state information, CF delivers reward information, PCs serve as the actor, and SCs serve as the critic. Triangle-headed arrows and circle-headed arrows represent excitatory synapses and inhibitory synapses, respectively. Dashed lines represent excitatory signals from CF via glutamate spillover (24).

In the cerebellar spiking network model, we implemented an RL algorithm known as the actor-critic model (18) based on the previous spiking actor-critic framework (21). In RL, an agent in state *s*(*t*) at time *t* takes action *a*(*t*) following a policy *π*. In response to action *a*(*t*), the environment changes its state and returns a reward *r*(*t*) to the agent. The agent learns to find the optimal policy that maximizes the expected future reward. Specifically, the agent of the actor-critic model consists of two modules: an actor and a critic. The actor calculates preferences of actions *h*_*α*_(*t*) and decides on action *a*(*t*), while the critic computes the state value *V* (*t*) which refers to the expected cumulative future reward. The temporal difference error (TD error) is calculated from the returned reward and the state value for updating the inner parameters of both the critic and the actor.

In our implementation, *s*(*t*) was transmitted through PFs to PCs, BCs, and SCs. The actor and the critic were PCs and SCs, respectively. Practically, the activity of PCs represented the avoidance of actions −*h*_*α*_(*t*), while the activity of SCs represented the additive inverse of the state value −*V* (*t*) rather than TD error which was represented by MLIs in a previous theoretical model (20). BCs contributed to action selection by modulating the activity of PCs (25, 26). Then, the most activated DCN group decided on an actual action *a*(*t*). In response to the action, reward information was transmitted through CFs to PCs and SCs.

#### Representation of states and rewards

States were represented by PFs following the formulation in the classical Marr-Albus-Ito model (1–3). For simplicity, we arranged the PFs on a two-dimensional grid plane composed of a number of small tiles (27). We named this plane the PF plane. Each tile had an individual receptive field that determined the PF’s firing rate based on the observed state, as described in the section titled ‘Implementations’.

In the present study, rewards *r*(*t*) were actually considered as punishments so that *r*(*t*) ≤0. CFs delivered the information as excitatory signals by flipping the sign as follows:

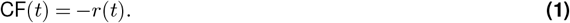

#### Critic

We designed the temporal activity of SCs to approximate the state value. Strictly speaking, the additive inverse of the state value was calculated by SCs as follows:

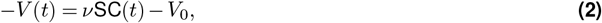

where *v*, SC(*t*), and *V*_0_ represent a scaling factor, the temporal activity of SCs, and the baseline of the state value, respectively. The baseline *V*_0_ was set at *ν* times the average value of SC(*t*), which was calculated during the latter half of an episode without learning. SC(*t*) was calculated as follows:

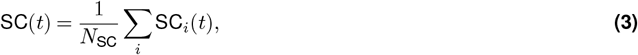

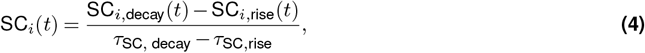

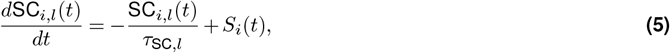

where *N*_SC_ and SC_*i*_(*t*) represent the number of SCs and the temporal activity of *i*th SC, respectively. *l* is a label in *{*rise, decay*}*. The temporal activity of *i*th SC rises with a time constant *τ*_SC,rise_ and decays to 0 with a time constant *τ*_SC,decay_. Spike train of the *i*th neuron is defined as 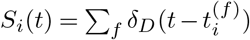 with the Dirac delta function *δ*_*D*_(*t*) and the *f* th spike of the *i*th neuron.

#### Actor and action selection

PCs performed the actor and determined the action taken (28). We interpreted the temporal activity of PCs as expressing avoidance of certain actions. Specifically, the PCs in our model compute the avoidance of an action 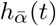 instead of the preference for an action *h*_*α*_(*t*). Therefore, the taken action *a*(*t*) was determined by the least-avoided action in action space 𝒜 at time *t*.

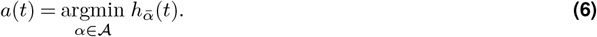

The avoidance of action *α* ∈ 𝒜 was represented by the activity of PCs in 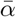 group as follows :

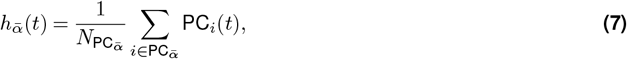

where 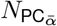 is the number of neurons in the group 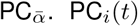 was the temporal activity of *i*th PC defined as follows:

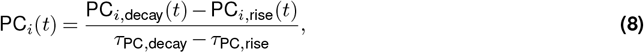

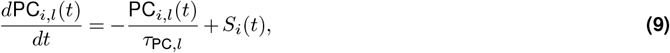

where *l, τ*_PC,*l*_, and *S*_*i*_(*t*) represent the label *l* ∈ {decay, rise}, the time constant for *l*, and the spike train of the *i*th PC, respectively. Practically, the actual action was generated at the downstream DCN (see the section titled ‘Implementations’).

Those spike activities of PCs were used for updating synaptic weights to realize which action was selected. It was vital that one PC group pauses while the other PC groups activate. We referred to this pausing behavior as a “dent”, while activation of neurons in a specific area was referred to as a “bump” by Frémaux et al. (21). The inhibitory loop between BCs and PCs helped to pause the single group solely to make a dent. Although the original spiking actor-critic framework supports continuous action space, the present model only supports discrete action space with only two actions, because of the difficulty of making a single dent on PCs.

#### Weight-update rules

Our weight-update rule was based on TD long-term potentiation (LTP) which had been proposed in a previous spiking actor-critic model (21). However, unlike the previous model, which calculated the time derivative of the state value directly to compute the TD error, our model did not calculate it, as there was no component in the molecular layer of our model that explicitly handled the TD error.

To address this problem, we conducted an eligibility trace and transformed the weight-update rule. As a result, our weight-update rule for the state value was as follows:

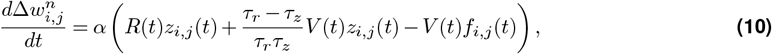

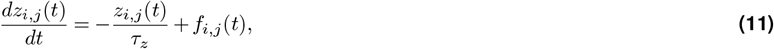

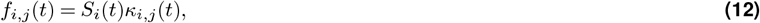

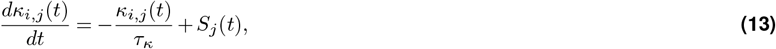

Where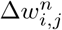, *α, R*(*t*), *z*_*i*,*j*_(*t*), *τ*_*r*_, *τ*_*z*_, *f*_*i*,*j*_(*t*), *S*_*i*_(*t*), *S*_*j*_(*t*), *κ*_*i*,*j*_(*t*), and *τ*_*κ*_ are the amount of synaptic change in *n*th episode, the learning rate, the negative reward, the eligibility trace for *w*_*i*,*j*_, the reward discount time constant, the decay time constant of eligibility trace, the forcing term of eligibility trace, the postsynaptic spike train, the presynaptic spike train, the window function for spike timing dependent plasticity, and the decay time constant of the window function. According to the previous model (21), the eligibility term memorizes the history of inputs before the time of the last postsynaptic neuron spike, *κ*_*i*,*j*_(*t*) is reset to 0 after the postsynaptic neuron spike. The relationship of our weight-update rule to the original weight-update rule (21, 29) is described in the section S1 of SI Appendix. The synaptic weight *w*_*i*,*j*_ was updated at the end of every episode *T*_end_ by 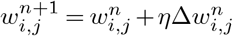 (*T*_end_) with learning rate *η*, and the synaptic weight was bounded between [0, 1]. Since SCs represented the additive inverse state value and we denoted the negative reward *r*(*t*) = −CF(*t*), the equation for the PF–SC synapses was expressed as follows:

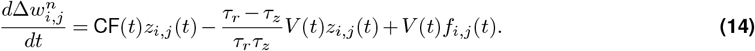

On the other hand, the weight-update rule for PF–PC synapses was expressed as follows:

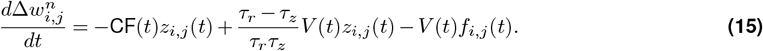

The specific learning rates for PF–SC synapses and PF–PC synapses are denoted as *η*_SC_ and *η*_PC_, respectively. Specifically, given that the conjunctive activation of PFs and CFs induces LTP at PF–SC synapses (30–33), our SCs learned the approximate additive inverse of the state-value. LTD at PF–SC synapses was due to PF firing, and was modulated by postsynaptic activation results in bidirectional synaptic plasticity (30–33). At PF–PC synapses, the conjunctive activation of PFs and CFs induces LTD (7), while PF activation induces LTP. These synaptic plasticities are also modulated by inhibitions of SCs (34), and we assumed that PC activity contributes to eligibility trace and bidirectional synaptic plasticity (35, 36).

Additionally, we confirmed that our critic is capable of learning state values through a Linear Track Task simulation (see SI Appendix, section S2), which is a simplified version of the water-maze task used to isolate learning by the critic from that by the actor (21).

### Simulation of Mountain Car Task

In order to evaluate the learning capability of our model as an RL machine, we applied our model to the mountain car task (37), a classic RL challenge, and evaluated its performance over 10 trials. In this task, an agent controls a car situated between two hills (Figure 2A). The goal is to have the car climb up the mountain to reach a flag positioned at the peak of the right hill. Due to the car’s limited engine power and the hill’s steepness, direct ascent is not possible. An agent observes a position *p*_*x*_(*t*) and a horizontal element of the car velocity *v*_*x*_(*t*), and decides the direction (either left or right) to push the car with a constant force 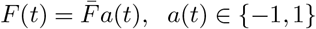. In early episodes, the car was confined to moving at the bottom of the valley, and failed to reach the goal (Figure 2B). However, after hundreds of episodes, our model learned to utilize the slope for acceleration, and eventually reached the goal (Figure 2C). Specifically, we found that the trajectory involved the car ascending the right hill, then the left, and finally the right hill again. The additive inverse of the state value −*V* (*t*), which was approximated by SC activity, was noisy but mostly stayed around 0 at every state before learning (Figure 2B). In a successful episode after learning (Figure 2C), the value was high at the start and decreased over time as the car approached the goal. The moving average of success rate over 10 trials progressively increased and stabilized at approximately 80% after 600 episodes (Figure 2D). These results indicate that our model solved the mountain car task, and suggest that our model successfully acted as an RL machine.

**Figure 2.**
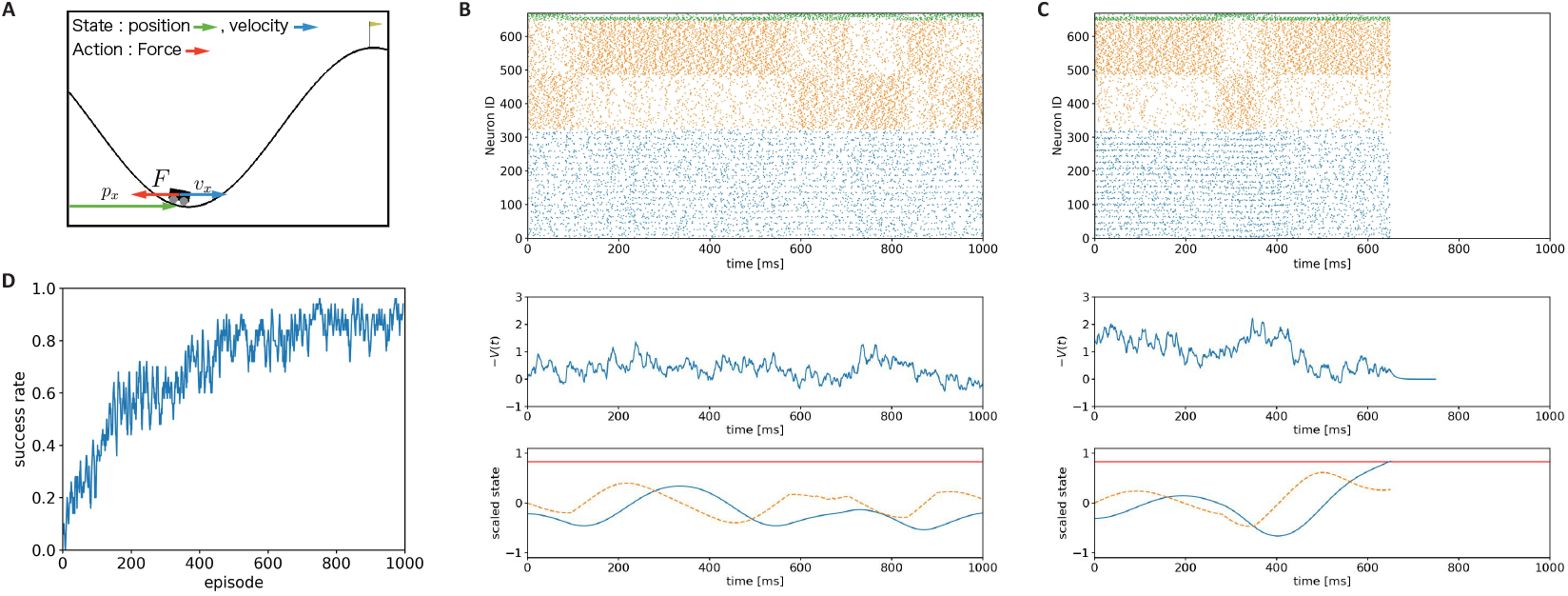
Simulation of the mountain car task. (**A**) Schematic of the mountain car task. *p*_*x*_ is the horizontal position of the cart, *v*_*x*_ is the horizontal velocity, and *F* is the external force applied to the cart by the agent. The flag marks the goal position. For clarity in the image, we adjusted the x-axis to start at 0 on the left edge. (**B**,**C**) Comparison of behaviors in the mountain car task. (**B**) Example behavior in early episodes. (**C**) Example behavior in later episodes. Top panels show raster plots of SCs (blue), BCs (orange) and PCs (green). The number of these neurons are 324, 324, and 20, respectively. Middle panels show trajectory of −*V* (*t*) computed from the activity of SCs. Bottom panels show trajectory of scaled states. Blue line and orange dashed line represent the normalized value of a position and that of a velocity. Red line represents the goal position. When the position reaches the goal, an episode ends while a simulation lasts 100 ms (see the section titled ‘Pre- and post-episode processes’). (**D**) Success rate of mountain car task. The moving average of success rate across 10 trials with window size 5 episodes.

### Simulation of Delay Eyeblink Conditioning Task

To examine whether our cerebellar model shows internal dynamics consistent with the biological cerebellum, we conducted simulation of a standard cerebellum-dependent motor learning task known as the delay eyeblink conditioning task (23). In this task, an animal is presented with two types of stimuli: a conditioned stimulus (CS) and an unconditioned stimulus (US) (Figure 3A). The CS, a neutral stimulus such as a tone, initially does not trigger any response. On the other hand, the US, such as an air puff, naturally induces an unconditioned blink reflex in the animal. However, when the CS is repeatedly paired with the US, the animal begins to anticipate the air puff whenever the CS is presented. This anticipation leads the animal to blink in response to the CS alone, a learned behavior known as the conditioned response (CR). We modified the task to have two-dimensional state space: time *t* and eyelid position *p*_*e*_(*t*), along with two actions: opening and closing the eyelid (Figure 3B). The time axis ranges from 0 ms to 1,000 ms, corresponding to the duration of stimulus presentation, while the eyelid position is measured on a scale from fully closed (0) to fully open (1). The CS was presented from 0 ms to 1,000 ms as a time signal, and the US was introduced at 500 ms as a punishment if the eyelid was open, i.e., if *p*_*e*_(500) was greater than 0.1.

**Figure 3.**
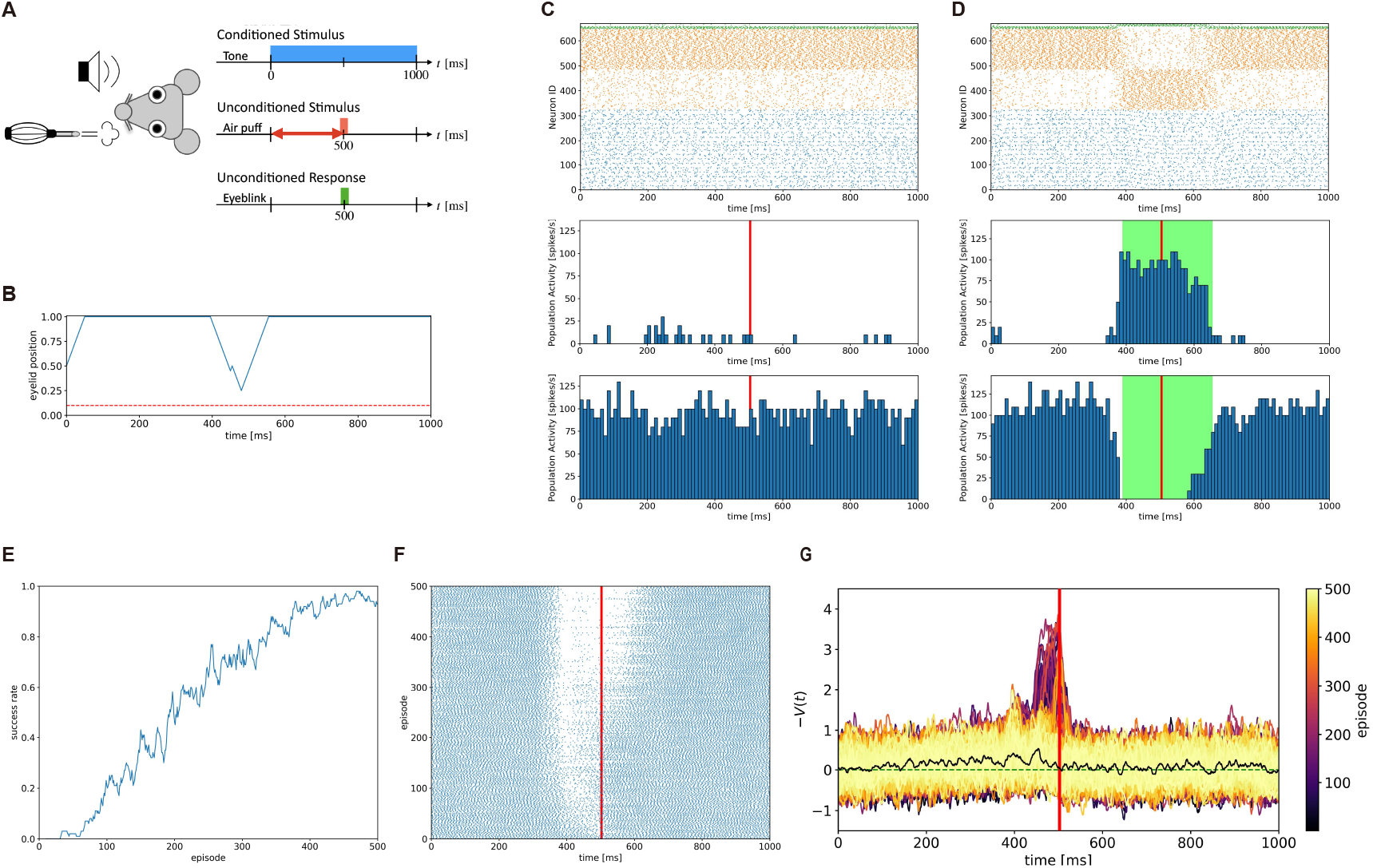
Simulation of the delay eyeblink conditioning task. (**A**) Schematic of the task. A neutral stimulus (CS) is presented, followed by an aversive stimulus (US) after a fixed interval. After repeated pairings, the agent closes its eyes (CR) in response to the CS alone. (**B**) Example state trajectory. The horizontal axis shows time from CS onset, and the vertical axis represents eyelid position. The dashed line marks the punishment threshold, triggered when the eyelid is above it at US onset (*t* = 500 ms). (**C**,**D**) Results for a failure episode (C) and a success episode (D). Top panels display SC (blue), BC (orange), and PC (green) raster plots. The number of these neurons are 324, 324, and 20, respectively. Middle and bottom panels show peristimulus time histograms for anti-open and anti-close PC population, respectively. Bin width is 10 ms. US onset is marked by a red line, and eyelid closure by green shading. (**E**) Moving average of success rate over 10 trials. (**F**) The activity changes of a single PC in the anti-close group, with dots representing spikes and red line indicating US onset. (**G**) Additive inverse of state value during conditioning, with colored traces showing stellate cell activity across 500 episodes (from dark purple in the 1st trial to light yellow in the 500th trial). The black line represents the average over the last 10 trials, green dashed line shows −*V* (*t*) = 0, and the red line indicates US onset.

In early episodes, the agent preferred to open the eyelid (Figure 3C and 3B), so the agent received the US as a negative reward. As learning progressed, the anti-open PC group became active, whereas the anti-close PC group paused as if PCs expected the US onset. Thus, the eyelid started to close before the US onset. For every 10 trials, the agent successfully acquired the CR to the CS: closing its eyelid to avoid a punishment before the arrival of the US (Figure 3E). Note that we defined a successful episode as a sufficiently closed eyelid to avoid the punishment at the US onset. When focusing on the activity changes of a single PC in the anti-close group (Figure 3F), the neuron acquired a behavior to pause before US onset. These behaviors were consistent with behavioral experiments of the delayed eyeblink conditioning task (23). We also analyzed the activity of SCs during the task (Figure 3G). In early episodes, since the agent received the punishment as the US, the SC activity as the additive inverse of the state value −*V* (*t*) increased exponentially toward the arrival of the US. These results are not inconsistent with biological experiments (38, 39), which show increases in MLIs’ firing frequency from the time of CS onset to the time of US onset. In later episodes when the agent successfully closed the eyelid more than 80% of the time, the SC activity returned to zero. This returning behavior resulted from the disappearance of CF signals due to the successful avoidance of punishments.

We also conducted a simulation of the same task without PF–MLI synaptic plasticity. The activity of the anti-close PC group decreased as it approached the US onset (Figure 4A). However, the agent was not able to elicit a stable closing response in our settings. In a later episode, the time to close the eye was delayed (Figure 4B) compared to the that in the intact simulation (Figure 3D), so the agent could not close the eye sufficiently. These results are also consistent with experimental results where MLI–PC inhibitions were impaired (40). Furthermore, these results suggested that our model successfully solved the eyeblink task with biologically plausible internal dynamics.

**Figure 4.**
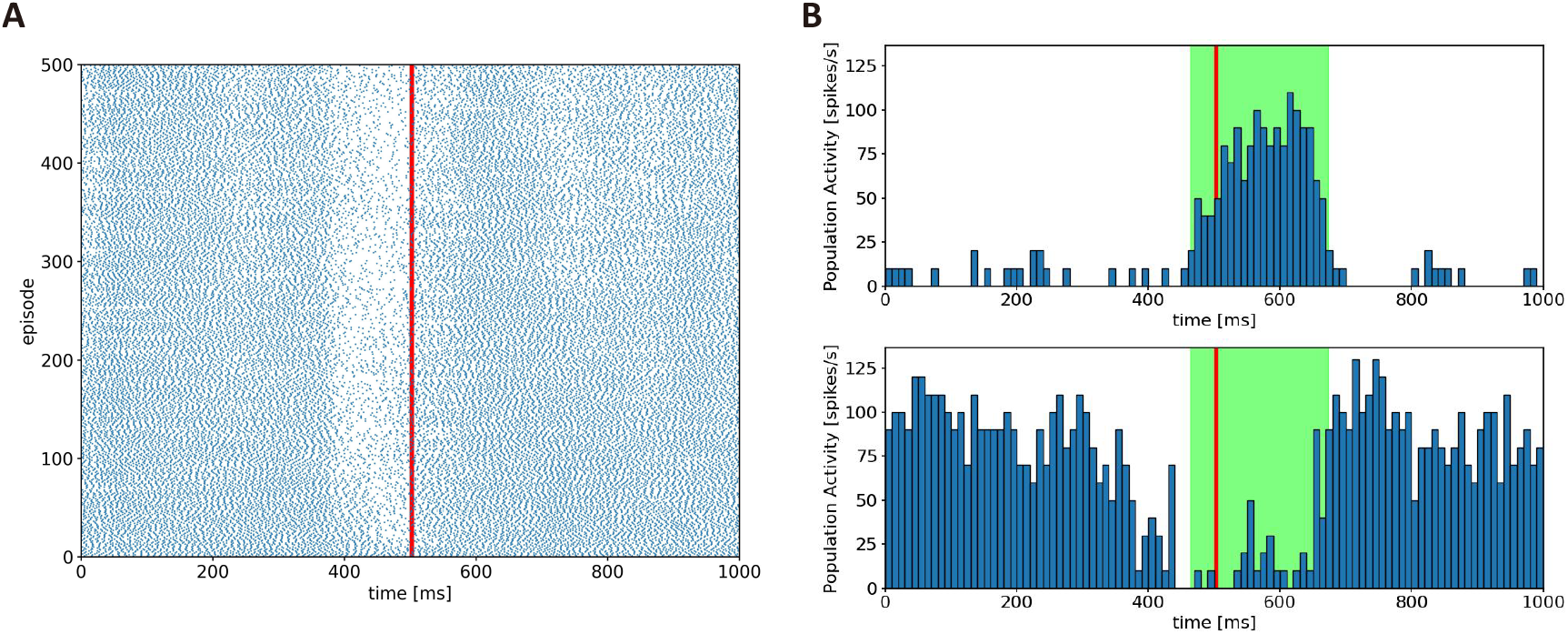
Results of the delay eyeblink conditioning task without PF–SC synaptic plasticity. (**A**) The activity changes of a single PC in the anti-close group. Dots represent spikes of the PC, and red line represents the time of US onset. (**B**) Example peristimulus time histograms of anti-open PC population (top) and those of anti-close PC population (bottom) in the later episode. Bin width is 10 ms. Red lines represent the time of US onset, while the green shaded intervals show when the agent closes its eyelid around the US onset.

## Discussion

In this study, we implemented the cerebellar spiking network model performing RL in an actor-critic manner. Our simulations confirmed the model’s ability to solve a simple RL task and demonstrated a cerebellum-dependent motor leaning task with biologically plausible internal dynamics. In contrast, Hausknecht et al. (13) reported that their cerebellar model was not able to solve RL tasks, only SL tasks. This is natural, because their model was formulated as an SL machine while ignoring PF–MLI synaptic plasticity. Recently, Geminiani et al. (17) implemented a spiking network-based cerebellar model that considers synaptic plasticity at both PF–PC and PF–MLI synapses. They reported that considering multiple synaptic plasticity mechanisms enhance learning capability in SL. Thus, those computational studies considered the cerebellum as an SL machine. Spiking network models of the cerebellum as an SL machine have been used for realtime motor control and online learning of hardware robots (14, 41). Our present cerebellar model will help those robotics studies by providing the capability of RL over SL.

In our cerebellar model, MLIs played essential roles as a critic. Recent studies have shown that MLIs actually contribute to learning in the cerebellum (38–40, 42) by their own learning capability through PF–MLI synaptic plasticity (30–32) and modulating PF–PC synaptic plasticity (34), in addition to controlling PC activities through MLI–PC inhibition (25, 26, 33, 43–46). In a behavioral experiment, genetically-modified mice lacking MLI–PC inhibition were able to acquire the CR, but the amplitude of eye closing was significantly lower than that of intact mice (40). Our results of the lesion simulation in the delay eyeblink conditioning task are consistent with the experimental results. These results suggest that the cerebellum can solve the association task even without normal SC activity, but the activity affects the performance of the acquired movement.

The present study assumed that CFs deliver negative reward information. In conventional cerebellar motor learning, CFs are considered to provide motor error signals, which are negative information. Furthermore, recent experimental results demonstrate that CFs carry not just motor-related information but also reward-related information such as reward delivery, prediction, and omission in mice (47–51). Thus, our assumption is consistent with those experimental findings. Hoang et al. (52) recapitulated the idea of negative reward information conveyed by CFs to Q-learning, which is a tabular-style reinforcement learning algorithm, and applied the Q-learning to a Go-No-go auditory discrimination task in mice. Furthermore, CF signals are provided in an all-or-none manner, and our model followed this property. However, several experiments have demonstrated that CF signals can carry graded information (53, 54), which might be useful to deliver reward information as analog values effectively.

In a standard RL algorithm called TD-learning, a reward-related signal tends to appear spontaneously at the onset of a signal that predicts the occurrence of a future reward after sufficient learning. The signal is called a reward-prediction signal or reward-prediction error. Ohmae and Medina (47) observed such reward-prediction signals conveyed by CFs at the CS onset after sufficient training of delay eyeblink conditioning in mice. Indeed, such prediction-error signals are prepared outside of the cerebellum. Therefore, in the present study, we did not consider such prediction error-type CF signals. If we incorporated them, those signals could have acted to control the balance between exploration and exploitation (18) while modulating the fluctuation of the internal network state. Another potential role would be to represent the most distant state from the goal state for reward delivery. Nevertheless, further investigation will be necessary to study the functional role of the prediction error-type CF signals.

If both the cerebellum and the basal ganglia perform RL, how do these regions cooperate for learning? One possible interpretation is hierarchical RL (HRL) (18) as discussed in our previous research (10). HRL considers two RL machines organized hierarchically. The higher RL machine breaks down a complex task into simpler, smaller sub-tasks. The lower RL machine tries to solve these subtasks one by one. Eventually, HRL involves creating a hierarchy of policies that makes the learning process more efficient and scalable (18, 55, 56). If the basal ganglia and the cerebellum cooperate to perform HRL, the resulting network would demonstrate powerful learning capabilities.

To our knowledge, our model is the first to implement cerebellar RL with spiking neurons. Our research provides an implementation that supports cerebellar RL theory to challenge the traditional view of the cerebellum as primarily a SL machine. Furthermore, our interpretation of a potential cooperative role of the basal ganglia and the cerebellum as HRL sheds light on how the multiple brain regions act together synergistically as a whole.

## Materials and Methods

### Implementations

First, we discretized a state space and mapped it on a PF plane with grid size *N*_*x*_ × *N*_*y*_. A tile positioned at (*i, j*) on the PF plane had its own responsible state **o**_*i*,*j*_ as follows:

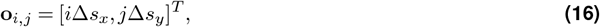

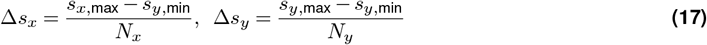

where *x* and *y* are the axes of the PF plane, and (*s*_*x*,min_, *s*_*x*,max_) and (*s*_*y*,min_, *s*_*y*,max_) represent the ranges of state *x* and *y*, respectively. When the agent observed state **o**(*t*) = [*x*(*t*), *y*(*t*)]^*T*^, PFs on a tile at (*i, j*) fired as an inhomogeneous Poisson spike generator with a firing rate *ρ*_*i*,*j*_(**o**(*t*)) defined as follows :

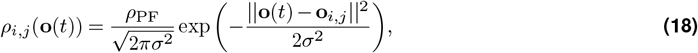

where *ρ*_PF_ = 50 Hz is a scaling factor of firing rate, and *σ* = 0.5 is a standard deviation. To maintain the total activity of all PFs when the agent is at the edge of a state space, we added “margin” tiles as an outer frame. Although we modeled PFs as Poisson spike generators for simplicity, more realistic implementations are reported (57).

We utilized a Leaky Integrate-and-Fire model (LIF) to SCs, BCs, and PCs. A LIF model was implemented as follows.

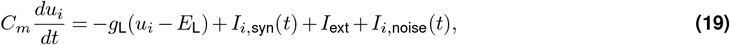

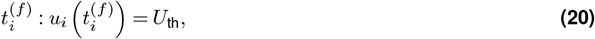

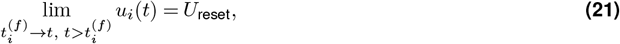

where *C*_*m*_, *u*_*i*_, *t, g*_L_, *E*_L_, *I*_*i*,syn_, *I*_ext_, and *I*_*i*,noise_ represent the membrane capacitance, the membrane potential of the *i*th neuron, time, the leak conductance, the resting potential, the synaptic current to the *i*th neuron, the external input current, and the noise current to the *i*th neuron, respectively. When the membrane potential of the *i*th neuron reaches the threshold potential *U*_th_, that time is denoted as 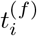, and the neuron fires the *f* th spike and resets its membrane potential to the reset potential *U*_reset_. A synaptic current was defined as follows:

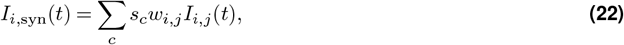

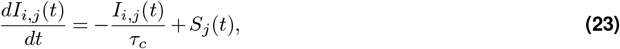

where *c* is a synapse label, *s*_*c*_, *w*_*i*,*j*_, *τ*_*c*_, and *S*_*j*_(*t*) are the scaling factor, synaptic weight between the *j*th presynaptic neuron and the *i*th postsynaptic neuron, decay time constant, and spike train of the *j*th presynaptic neuron, respectively. Whether it is an inhibitory or excitatory current is determined by the sign of the scaling factor. Noise current *I*_*i*,noise_(*t*) was modeled as follows:

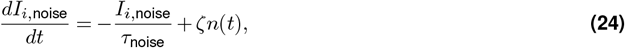

where *τ*_noise_ is a time constant, and ζ is an amplitude factor. *n*(*t*) is a uniform random variable in the range [ −1, 1]. Parameters of all neurons were set as listed in Table 1, and parameters of all synapses were set as listed in Table 2.

**Table 1.** Neuron-specific parameters. These parameters were adjusted from those used in our previous model (15).

| Type | $C_m$ [pF] | $g_L$ [mS] | $E_L$ [mV] | $U_{th}$ [mV] | $U_r$ [mV] | $I_{ext}$ [pA] | $\zeta$ [pA] |
| --- | --- | --- | --- | --- | --- | --- | --- |
| SC | 107 | 2.32 | -68.0 | -55.0 | -70.0 | 30.0 | 5.0 |
| BC | 107 | 2.32 | -68.0 | -55.0 | -70.0 | 30.0 | 1.0 |
| PC | 107 | 2.32 | -65.0 | -55.0 | -66.0 | 30.0 | 5.0 |

**Table 2.**
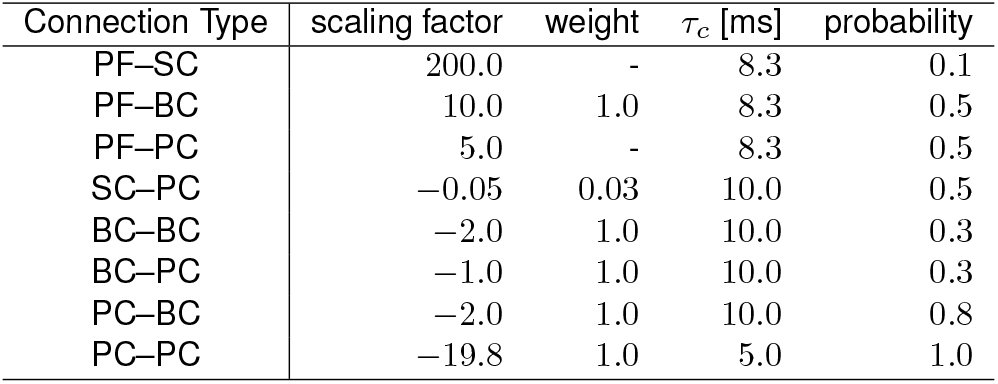
Synaptic parameters for each connection type. A hyphen in a weight cell means it will be modified because of synaptic plasticity.

As mentioned above, PFs were distributed on the two-dimensional grid plane, and the grid size was determined by each task. Each tile contained 100 PFs. SCs were prepared 1 per tile. Each SC positioned at (*i*Δ*s*_*x*_, *j*Δ*s*_*y*_) could receive excitatory inputs from PFs located in *s*_*x*,min_ ≤*x* ≤*s*_*x*,max_ and (*j* − 1)Δ*s*_*y*_ ≤*y* ≤ (*j* + 1)Δ*s*_*y*_. The number of BCs was the same as that of SCs, and the number of PCs was 20. Both BCs and PCs could receive excitatory PF inputs from all tiles. The BC–BC, BC–PC, PC–BC, and PC–PC connections were constructed based on the grouping manner mentioned in the section titled ‘Spike-based implementation of cerebellum-style RL’. The actual synapses were formed stochastically (Table 2). In this study, we divided BCs and PCs into two groups for all tasks. Additionally, for the sake of simplicity, we assumed that all CFs synchronized their activity so that our model had a single CF. Synaptic plasticity was applied to PF–SC and PF–PC synapses. The details of the weight-update rules are described in Equations Eq. (14) and Eq. (15).

Finally, the outcome of actions were generated by DCN cells prepared in an equal number of PCs. Each DCN cell was inhibited by its counterpart PC. This configuration means that the activity of each DCN cell, denoted as DCN_*i*_, was modulated by the corresponding PC activity PC_*i*_(*t*) defined by Equation 8, as follows:

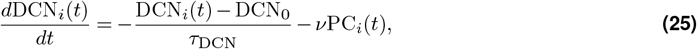

where *τ*_DCN_, DCN_0_, and *ν* represent the decay time constant, baseline activity of DCN, and a scaling factor, respectively.

Since PCs in the group 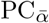 represented the avoidance of action *α* and inhibited DCN cells, the activity of DCN group DCN_*α*_ represented the preference of actions *h*_*α*_(*t*) as follows:

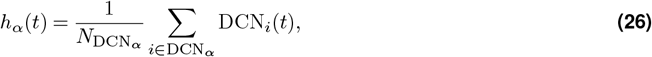

where 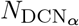 is the number of neurons in the group DCN_*α*_, and DCN_*i*_ (*t*) is the activity of *i*th DCN in the group DCN_*α*_ corresponding to action *α*. Therefore, the most activated DCN group represented the most preferred action so that the final outcome of actions was chosen as follows:

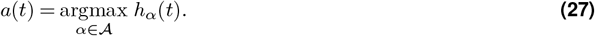

### Task Definitions

#### Pre- and post-episode processes

As a pre-episode process, at the start of each episode, a free run for 200 ms was carried out to discard initial transient activity that could affect learning. Then, at the end of each episode, a simulation continued for 100 ms while stopping the firing of PFs to update synaptic weights. These pre- and post-episode processes were performed in all tasks.

#### Mountain Car Task

The dynamics of the car are described by the following differential equations :

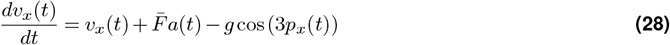

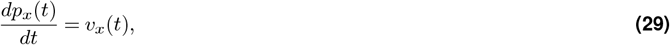

where 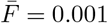 is a force amplitude, *a*(*t*) ∈ {−1, 1} represents an action as the force direction, *g* = 0.0025 represents the gravity. Observation spaces of the velocity and position are bounded by [−1.2, 0.6] and [−0.07, 0.07], respectively. When the car collides with the wall, its velocity is immediately reset to 0 without any negative reward associated with the collision. An agent receives constant negative reward at every step in the original settings. However, the maximum firing frequency of CF, which is approximately 10 Hz (58), was not enough to represent the constant negative reward. Therefore, in this study, we modeled a new negative reward function for CF. CF fired when the car was far from the goal and the speed was slow. Thus, the firing rate *v*(*t*) was defined as follows:

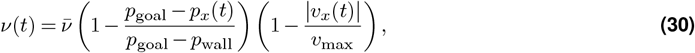

where 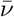, *p*_goal_, *p*_wall_, and *v*_max_ are max firing rate of 5 Hz, goal position, left side wall position, and limit of speed respectively. Then a negative reward function is defined as follows:

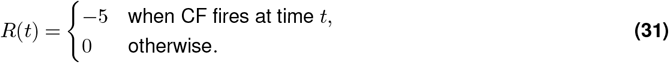

The car starts from the bottom of the valley with a velocity of 0 in every episode. The action *a*(*t*) represents the direction in which the agent pushes the car. An episode ends when the car reaches the goal or when the elapsed time exceeds 1,000 ms, with no special negative reward given.

The grid size of the PF plane was 16 × 16. The parameters of the state value function, as defined in Equation Eq. (2), were *ν* = 200 and *V* _0_ = 2.48. The reward discount time constant *τ*_*r*_, the decay time constant of eligibility trace *τ*_*z*_, and the decay time constant of the window function *τ*_*κ*_ were set to 100 ms, 20 ms, and 20 ms, respectively. Initial synaptic weights of PF–SC and PF–PC were 0.05 and 0.8, respectively. Both learning rates for the critic and the agent were 0.4.

We conducted 10 trials of 1,000 episodes each. We calculated the changes in success rate for each trial using a moving average of results (success=1, failure=0) with a window size of 10 episodes, and then calculated the average across trials.

#### Delay Eyeblink Conditioning Task

The dynamics of the eyelid is described as follows:

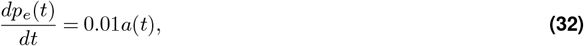

where *a*(*t*) ∈ {−1, 1} represents the action the agent takes at time *t*. Also, negative reward is defined as follows:

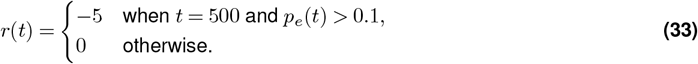

The grid size of the PF plane was 16 × 16. The parameters of the state value function, which defined in Equation Eq. (2), were *ν*= 200 and *V*_0_ = 2.48. The reward discount time constant *τ*_*r*_, the decay time constant of eligibility trace *τ*_*z*_, and the decay time constant of the window function *τ*_*κ*_ were set to 100 ms, 20 ms, and 20 ms, respectively. To prioritize the ‘open’ action, we initially set synaptic weights for PF–PC_anti close_ to 0.8, while PF–PC_anti open_ was set to 0.7. We conducted the simulation for 500 episodes with learning rates *η*_SC_ = 0.05 and *η*_PC_ = 0.3.

The whole code of our model and all environments were written in C++. All ordinary differential equations were solved numerically with a forward Euler method with a temporal resolution of 1.0 ms.

## Author contributions

T.Y. conceived and designed the research. R.K. and H.Y. contributed the formulation and the implementation of the algorithms. R.K. wrote the code, performed simulation, and analyzed the data. R.K. and T.Y. wrote the original draft. R.K., H.Y., and T.Y. discussed the draft and revised it. All authors contributed to the article and approved the submitted version.

## Acknowledgment

We would like to thank Profs. Kazuo Kitamura, Jun Igarashi, Shogo Ohmae, and Kenji Doya for their fruitful discussions and helpful insights.

## Supplementary Information

### A. Relationship between our weight-update rule and the previous weight-update rule

The weight-update rule and eligibility trace in our model are defined as follows:

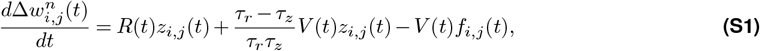

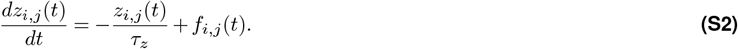

By incorporating the definitions, 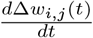 can be transformed as follows (Equations):

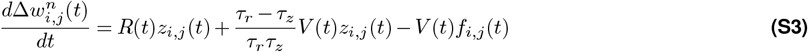

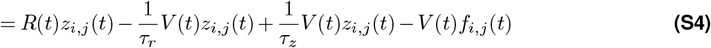

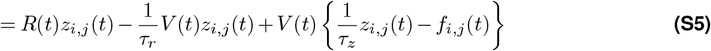

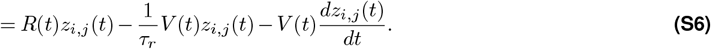

The total weight change over an episode Δ*w*_*i*,*j*_(*T*_end_) is expressed as follows:

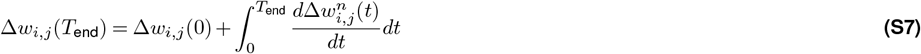

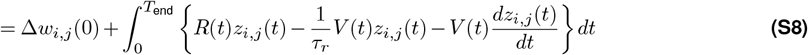

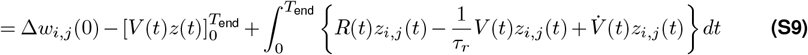

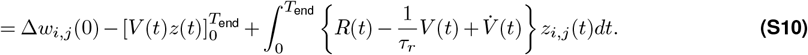

Here, by assuming Δ*w*_*i*,*j*_(0) = 0, *z*(0) = 0, and *V* (*T*_end_) = 0, the following equation is obtained:

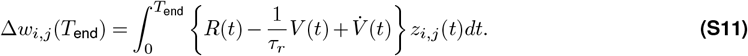

The expression within the curly brackets is equivalent to the continuous time TD error defined by (1), which was used by the previous spiking actor-critic framework (2).

### B. Simulation of Linear Track Task

In order to evaluate whether our critic could approximate −*V* (*t*), we conducted a simulation of the linear track task (2). In this task, the environment is a narrow rectangular plane (Figure S1A), on which the agent consistently moves from the leftmost side (start) to the rightmost side (goal) at a fixed speed. Thus, the agent always reaches the goal at the same time, and receives the same negative reward across all episodes. Once the time constant of the reward discount *τ*_*r*_ is determined, the theoretical value can be solved (Figure S1A).

In this task, the plane size was 40 × 1, and a fixed starting position was *s* = 5. The agent ran with a fixed speed of 0.05 per ms toward a goal area (*s >* 39). When the agent reached the goal, a negative reward of −5 was given. The parameters of the state value function, which defined in Equation 2, were *ν* = 300 and *V*_0_ = 12.4. The learning rate and the initial weight of PF–SC were 0.04, and 0.05. The reward discount time constant *τ*_*r*_, the decay time constant of eligibility trace *τ*_*z*_, and the decay time constant of the window function *τ*_*κ*_ were set to 100 ms, 20 ms, and 20 ms, respectively.

On the first run, the SC represented a noisy, around zero value function (Figure S1B). Over 100 trials, the SCs learned the value function progressively. Then the SC activity increased exponentially toward the goal, as if it expected the given negative reward (Figure S1B). The averaged value in 20 late trials, as plotted with a black line, nicely matches the theoretical value function. These results were consistent with previous research (2), and suggest that our weight-update rules were successful and that the SCs were able to approximate the additive inverse of the state value −*V* (*t*).

**Figure S1.**
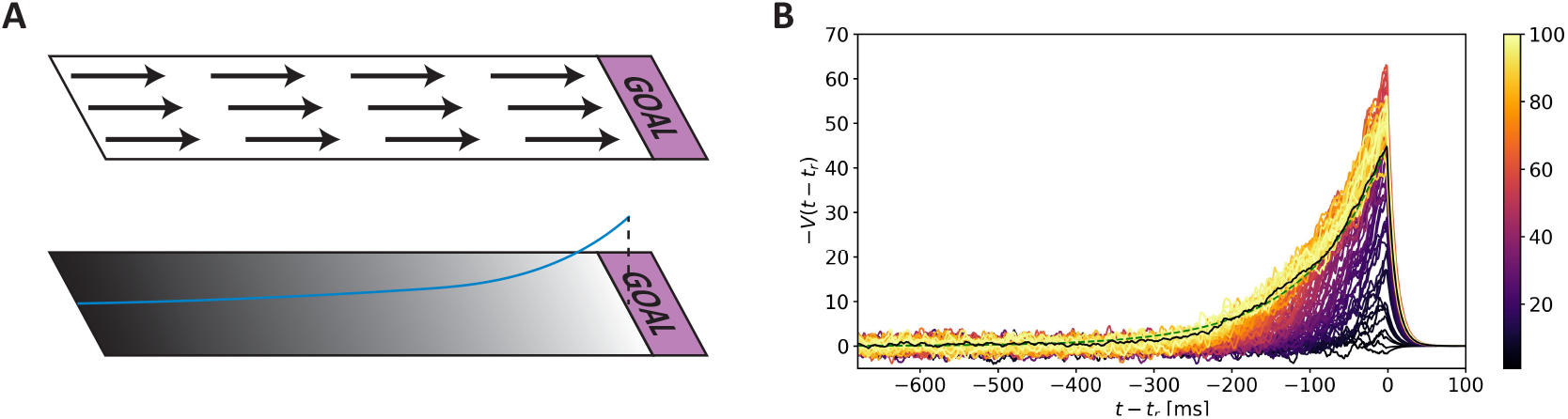
Simulation of Linear Track Task. (**A**) Environment of the linear track task. Top panel shows the environment and fixed policy. The agent starts at the left edge, and moves straight to the goal shown as a pink area. Bottom panel shows theoretical state value. When the agent follows a fixed policy, the theoretical state value is uniquely determined. (**B**) Additive inverse of the state value learned by the critic. Each colored trace shows the value represented by the stellate cell neurons activity against time in 100 first simulation trials (from dark purple in the 1st trial to light yellow in the 500th trial). The plots are aligned with the timing of the negative reward delivery *t*_*r*_. The black line shows the average of over last 10 trials. The green dashed line shows the theoretical value function.

